# A Novel Approach to Topological Network Analysis for the Identification of Metrics and Signatures in Non-Small Cell Lung Cancer

**DOI:** 10.1101/2022.11.22.517587

**Authors:** Isabella Wu, Xin Wang

## Abstract

Non-small cell lung cancer (NSCLC), the primary histological form of lung cancer, accounts for about 25% - the highest - of all cancer deaths. As NSCLC is often undetected until symptoms appear in the late stages, it is imperative to discover more effective tumor-associated biomarkers for early diagnosis. Topological data analysis is one of the most powerful methodologies applicable to biological networks. However, current studies fail to consider the biological significance of their quantitative methods and utilize popular scoring metrics without verification, leading to low performance. To extract meaningful insights from genomic data, it is essential to understand the relationship between geometric correlations and biological function mechanisms. Through bioinformatics and network analyses, we propose a novel composite selection index, the C-Index, that best captures significant pathways and interactions in gene networks to identify biomarkers with the highest efficiency and accuracy. Furthermore, we establish a 4-gene biomarker signature that serves as a promising therapeutic target for NSCLC and personalized medicine. We designed a Cascading machine learning model to validate both the C-Index and the biomarkers discovered. The methodology proposed for finding top metrics can be applied to effectively select biomarkers and early diagnose many diseases, revolutionizing the approach to topological network research for all cancers.

## Main

Lung cancer is the deadliest cancer worldwide and is highly fatal, with a five-year survival rate of <21%^1^. Non-small cell lung cancer (NSCLC) accounts for about 85% of all lung cancer cases^2^. More than half of lung cancer patients die within one year of being diagnosed because NSCLC is often not detected until the late stages with the onset of noticeable symptoms^3^, making treatment extremely difficult. This clinical need has driven the search for more effective prevention and early diagnosis strategies, including the identification of effective biomarkers.

By providing insights into the molecular origins and behaviors of NSCLC, biomarkers help identify high-risk patients and targets for personalized medicine and the development of targeted therapies^4,5^. The expansion of the availability and quantity of molecular biological data has created a pressing need for improved computational methods for data analysis. In the past few years, topological data analysis (TDA) has revolutionized the oncology field, becoming one of the most powerful and widespread tools to extract useful information from high-dimensional biomedical data^6,7^. We can use TDA to analyze the interactions between genes and identify essential biomarkers. Network nodes are ranked by scoring methods that reflect varying network features^8,9^. The high-scoring nodes are selected as the most essential genes in the network and further analyzed as potential markers.

Despite recent advances in applying TDA to cancer studies, one critical issue remains largely overlooked – whether geometric and topological connectivity implies functional connectivity in biological networks^10,11^. A comprehensive analysis of the correlation between geometric connectivity and functional connectivity is currently lacking, leading to ineffective use of the powerful methodology. The performance of existing works is not high, largely due to the lack of understanding of the most key aspect of topological analysis – the network scoring metrics. They rank the network nodes by different features, evaluating the network in varying aspects to select the high score nodes^11^. Inherently, these scoring metrics are defined geometrically with no clear implications for functional significance. Existing studies often fail to biologically validate the scoring methods they choose to use in biomarker identification for disease prediction^12,13^. They often utilize popular and conventional geometric-based scoring metrics (e.g. Degree)^14–16^ without reasoning, and use only a single scoring metric.

Thus, the central goal of this study is to identify the top-performing topological scoring metric (or method) that best captures functional significance in protein networks. More specifically, we focus on designing a metric, or a composition of metrics, that identifies critical biomarkers most efficiently and accurately. To do so, we conducted systematic and comprehensive analysis and validation in two main stages.

First, to investigate the ability of the metrics to select effective biomarkers, we performed detailed functional enrichment and network analyses to identify the top biomarkers for NSCLC diagnosis. To exploit the power of gene interactions, we further explore the concurrent use of multiple biomarkers in NSCLC prediction and develop a biomarker-signature consisting of a small group of top biomarkers. To determine their diagnostic ability, we introduce a method to calculate the area under the receiver operating characteristic curve (AUC) for multiple biomarkers concurrently, known as Integrated AUC. By doing so, we directly evaluate the biomarkers by their functional abilities to diagnose NSCLC. In the second stage, we build off the previous stage to identify the top-performing topological scoring metrics (or methods) by evaluating their ability to select biomarkers with the highest diagnostic capabilities most efficiently and accurately. We propose a novel composite selection index that concurrently considers complementary factors to evaluate the network and produce biologically significant results.

We expect the proposed methods to fundamentally advance topological network research in the cancer field. We unlock a better understanding of how geometric relationships in gene networks relate to functional mechanisms to extract meaningful insights from genomic data. Our proposed methods are not restricted to the use in diagnosing NSCLC, but can be extended to the early diagnosis of other types of cancers. With the widespread use of topological network analysis, we expect the proposed techniques to drastically improve the search for new biomarkers and targets for drug therapy and redefine the usage of TDA in oncology.

## Results

### Biomarker signature identification

Identifying specific and sensitive biomarkers is critical for early cancer diagnosis^17^. As the first step of this study, we identified the top biomarkers to serve as a base for us to explore in depth the critical scoring metrics for biomarker identification in the next stage.

#### DEG screening and functional enrichment

In total, we identified 267 differentially expressed genes (DEGs) (p-value<0.01 and |logFC| > 1.2) from the three datasets GSE31210, GSE33356, and GSE50081, taken from the Gene Expression Omnibus (GEO)^18^. 93 are upregulated and 174 are downregulated (Fig.1a). To investigate the roles that the DEGs play in disease mechanisms, we must the investigate DEG-related pathways. Through enrichment analyses, we identified 10 enriched GO terms with FDR < 0.05 (Fig.1c), and their z-score expression values (Fig.1b.)

**Figure 1.**
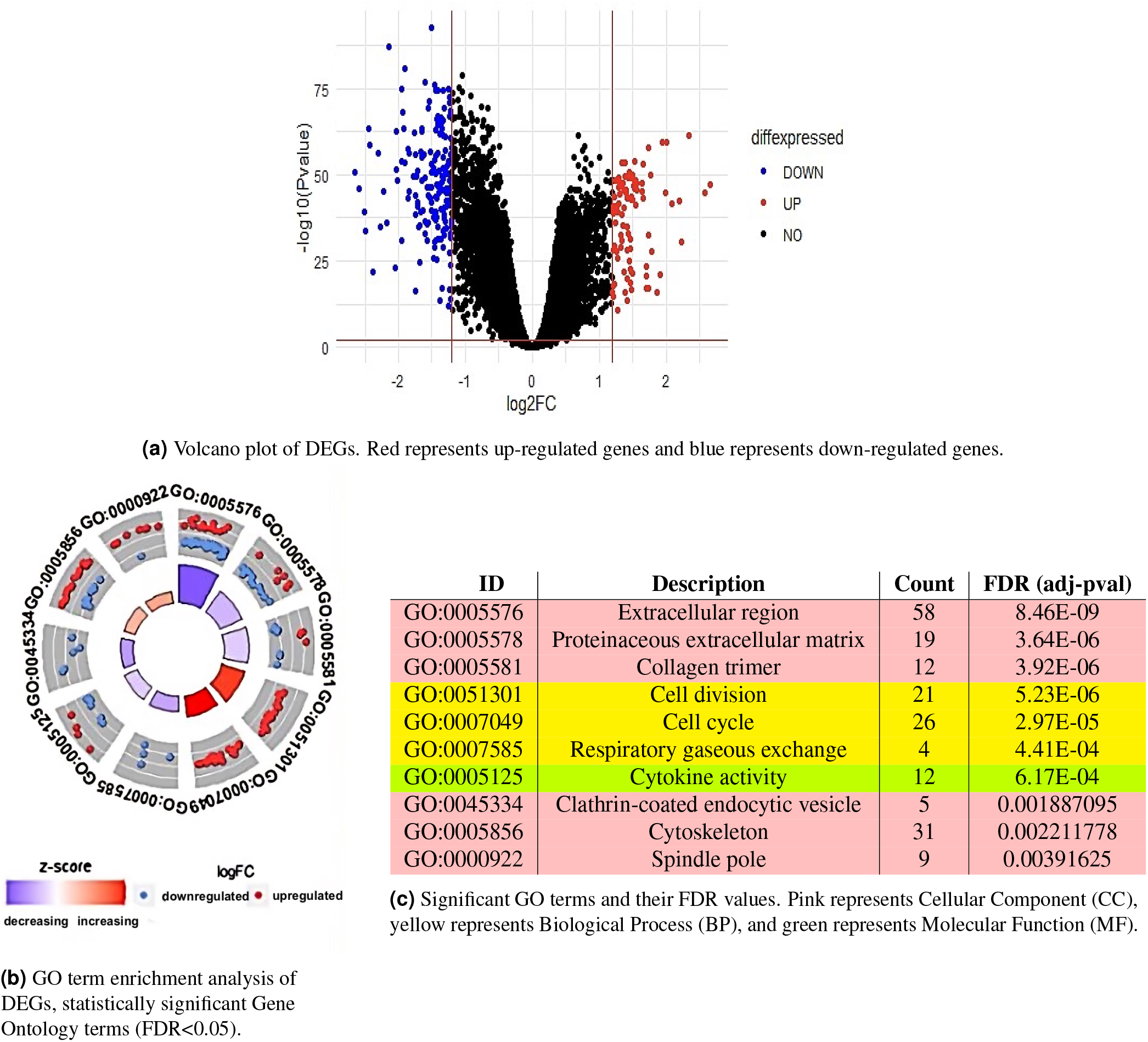
GO and KEGG analyses of DEGs

Upregulated DEGs play a role in multiple pathways that promote tumorigenesis, including cell division (GO:0051301), cell cycle (GO:0007049), cytoskeleton (GO:0005856), and spindle pole (GO:0000922). Downregulated genes were significantly involved in weakened tumor defense and disruptions in signal transduction pathways, such as decreased cytokine activity (GO:0005125) and clathrin-coated endocytic vesicles (GO:0045334). Downregulated genes were also enriched in the extracellular matrix (GO:0005576 and GO:0005578), and ECM-receptor interaction (hsa04512, FDR= 0.0152) was identified through KEGG pathway analysis. Complications in the ECM-receptor interaction pathway can result in induced cancer progression and development, as ECM-receptors play important roles in tumor shedding, adhesion, and degradation^19^. Our results show that the DEGs are largely connected to disease-related pathways and play potent roles in cancer onset and development.

#### PPI network analysis

Disease susceptibility and other disease correlated factors are due to the perturbation of an interconnected gene network^20^, not single gene mutations in isolation. To explore the interactions between DEGs and identify intrinsic mechanisms of disease, we constructed a protein-protein interaction (PPI) network (Fig.2) based on our 267 DEGs to understand the topology of molecular interactions and identify the most essential top-scoring biomarkers in the network. The network was visualized using Cytoscape^21^ and further analyzed with the CytoHubba algorithm^22^. The topological scoring metrics in CytoHubba are divided into two categories, *local* to evaluate individual nodes and *global* to evaluate the network as a whole. The local metrics include Degree, Maximal Clique Centrality (MCC), Density of Maximum Neighborhood Component (DMNC), Maximum Neighborhood Component (MNC), and Clustering Coefficient. The global metrics include Betweenness, Bottleneck, Eccentricity, Closeness, Radiality, Stress, and Edge Percolated Component (EPC). Often, literature studies utilize only one scoring method^14–16^. To ensure that no essential genes are missed and all possibilities are considered, we created a complete and comprehensive list of candidate biomarkers using all twelve scoring methods.

**Figure 2.**
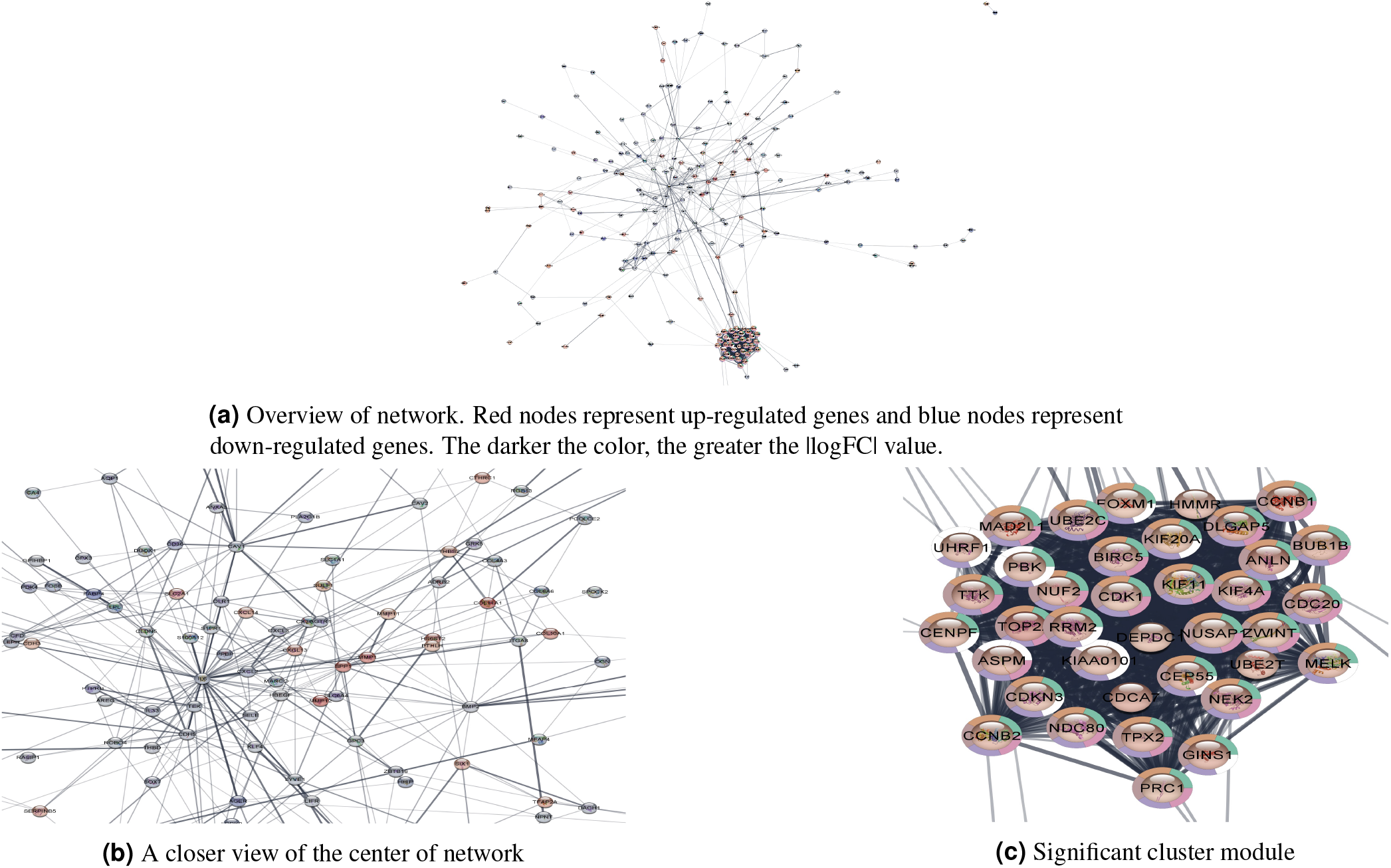
PPI network of DEGs and CytoHubba visualization

Each metric was utilized to select 10 top nodes each, with some having a higher cutoff because a few nodes share the same ranking score. Without counting overlapping genes between metrics, we obtained 82 candidate biomarkers (Table 1a) overall. To evaluate the ability of biomarkers in distinguishing between disease and control, we chose to use area under the receiver operating characteristic curve (AUC) score to select the overall top 20 disease-correlated genes in the network (Table 1b). This allows us to directly evaluate the biomarkers by their ability to predict disease.

**Table 1.**
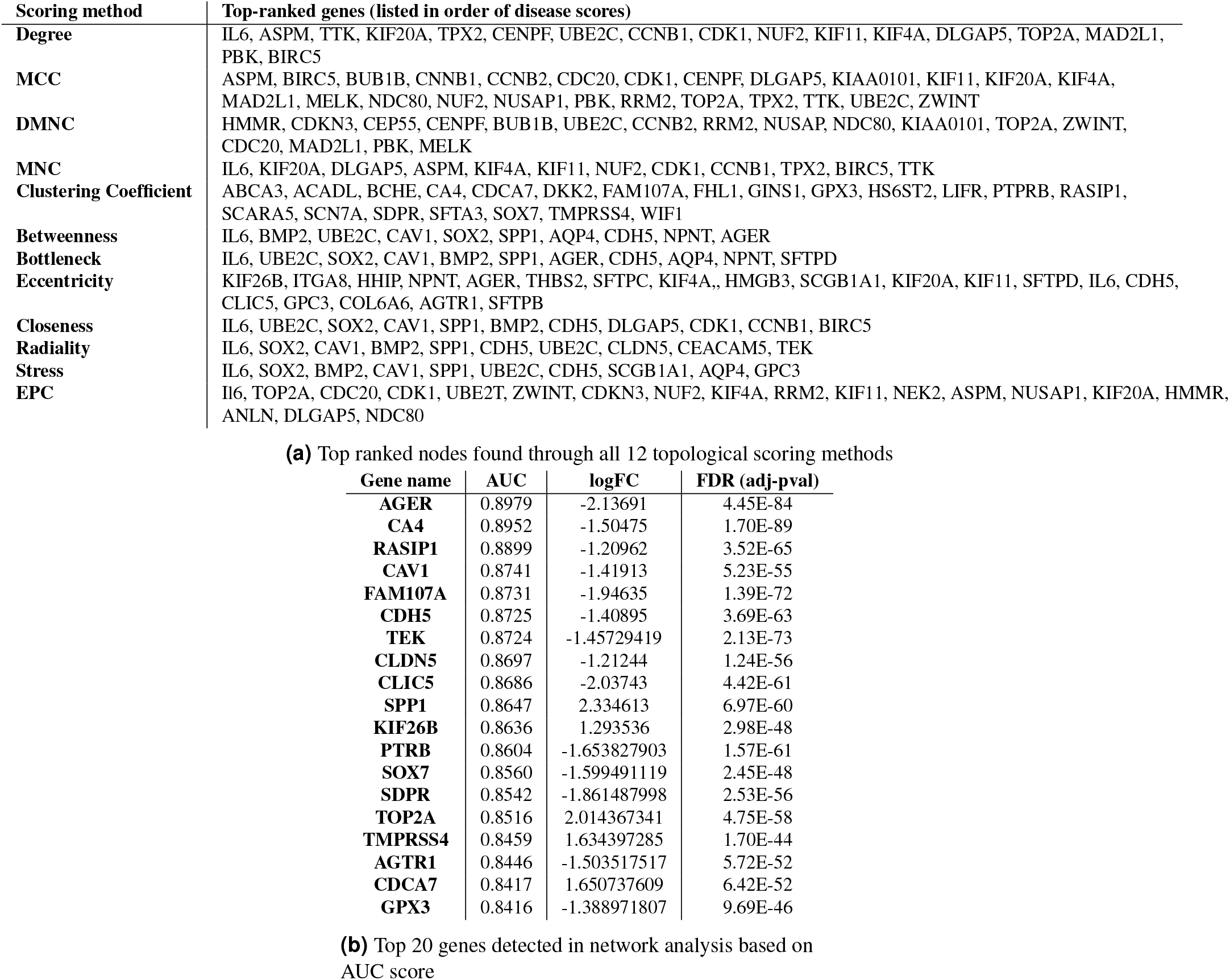
Top biomarkers in the PPI network based on all 12 network scoring metrics

#### Disease prediction with multiple biomarkers simultaneously

Cancer is caused by multiple genes in a functional or signaling pathway working together in a cascade of mutations to promote tumorigenesis, and can never be caused or predicted by a single mutation or gene. Rather than only considering the capability of individual biomarkers as commonly done in literature studies, we further explore the concurrent use of multiple biomarkers in NSCLC prediction to vastly increase the diagnostic performance, Utilizing multiple biomarkers is more comprehensive, and may better deal with disease heterogeneity and reduce anomalies during prediction. The incremental usefulness of adding multiple biomarkers from different disease pathways has not, to our knowledge, been fully evaluated amongst other NSCLC studies. Simply using too many may decrease performance. We need to find the optimal number of biomarkers to use concurrently for the highest performance, as well as a way to evaluate the joint performance of biomarkers.

As each biomarker’s expression value quantifies its relationship to the health condition of a subject, we propose the concept of *Integrated AUC* to calculate the AUC of the aggregated expression of biomarkers. In this study, the aggregated expression is defined as the mean expression of the group of biomarkers as it is most suitable, but its definition may be expanded.

We found that the top 4 biomarkers *AGER, CA4, RASIP1*, and *CAV1* produced the highest Integrated AUC at 0.9238 (Fig. 3a). They make up our **4-gene biomarker signature** for the prediction of NSCLC. Their Receiver Operating Characteristic (ROC) curves are visualized in Figure 3c. For comparison purposes, we also calculated the AUC of the top 10 genes combined. As expected, the 4-gene signature outperformed 10 genes (Fig. 3b). This may be due to the nature of interactions between genes – certain biomarkers may not interact optimally for high performance, or biomarkers outside of the top four may play roles of lower significance. The finding of the biomarker signature and the use of Integrated AUC as an evaluation metric can be expanded on and further explored to improve the prediction accuracy of other types of cancers.

**Figure 3.**
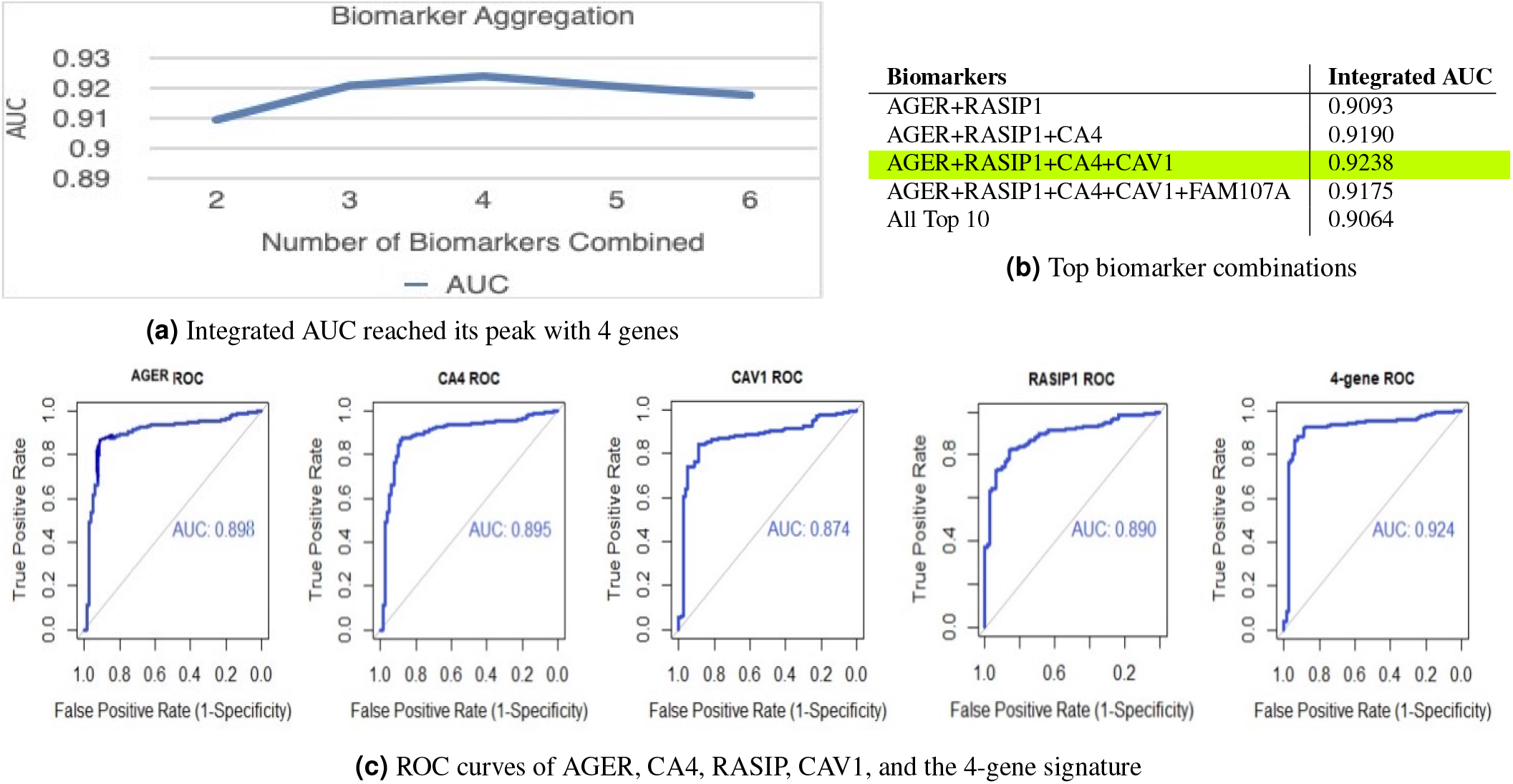
Performance of 4-gene signature

#### Validation of 4-gene signature by survival analysis and TCGA database

At the beginning of the study, we divided the dataset into 80% for identification of biomarkers and metrics, and 20% validation. To validate the effectiveness of using the 4-gene signature, we compared its performance in the validation data set to each of its 4 individual components, as well as that of the top 10 genes combined. The 4-gene signature outperformed all of its individual genes, as well as the top 10 genes together (Supplementary Table 1), demonstrating that it is less complex and more effective than using more genes, and more powerful than only using individual biomarkers alone. To validate the effect of the 4-gene signature on NSCLC prognosis, we performed overall survival analysis with these genes using Kaplan-Meier survival plots (Fig. 4a) to examine their impact on patient survival, with a threshold of p-value < 0.01 to determine significance. The low expression of *AGER, CA4, RASIP1*, and *CAV1* are all associated with poor overall survival, indicating their significant roles in NSCLC prognosis.

**Figure 4.**
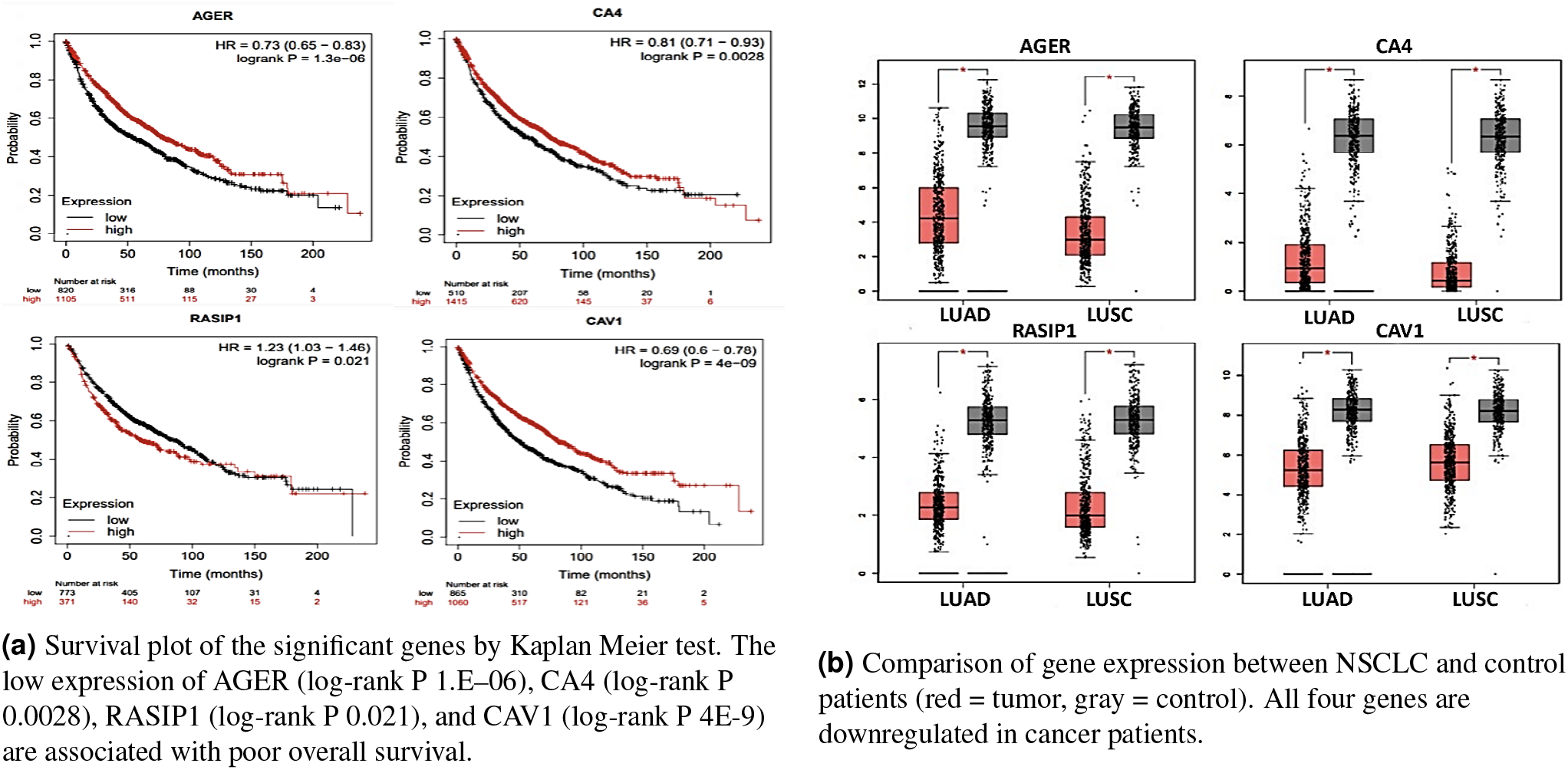
Differential expression of 4-gene signature in NSCLC patients

To confirm that our results are applicable outside our data set, TCGA data from the GEPIA interactive website was used to verify the identified genes to be effective amongst other NSCLC cases. Figure 4b compares gene expressions from two histological types of NSCLC, lung squamous cell carcinoma (LUSC) and lung adenocarcinoma (LUAD), and normal lung tissues. All four genes are significantly downregulated in cancerous patients, indicating that the four-gene signature can be expanded and used for new patients and data. From our GO enrichment analysis, these genes play important roles in the ECM matrix and signalling pathways. They are important for cancer detection and treatment, and can act as therapeutic targets for future drug therapy and personalized medicine.

### Critical Network Topological Metrics

In the past decade, topological data analysis has grown into a prominent role in the oncology field. The scoring metrics are crucial to topological data analysis as they identify the most influential network nodes. Scoring metrics, however, describe only geometric relationships between nodes in the network, not their relation to disease diagnosis. Consequentially, the significance and implications of the scoring methods are vastly overlooked in most biological studies, and a single metric is often employed without reasoning. The most frequently used metric is Degree, a local metric that may not capture the full extent of gene network interactions. Our foremost goal is to thoroughly analyze the performance of these quantitative methods applied to biological networks and identify the top metrics that best capture functional connectivity and biological implications.

We set out to achieve this in two major stages. We first evaluate all 12 topological scoring metrics based on their diagnostic ability for effective biomarker selection. In the search for biomarkers, using a single metric may be inadequate, while using all metrics together is complex and inefficient, and possibly erroneous. Therefore, in the second stage, we further investigate the performance of using multiple metrics concurrently in diagnosis and design a powerful metric. Few previous studies have meaningfully utilized multiple network metrics concurrently to look for biomarkers. Our investigations indicate that increasing the analytical coverage of a biological system through optimal pairings of metrics leads to more robust results.

#### Evaluating the performance of individual metrics

To evaluate the ability of the metrics to identify essential biomarkers, we calculated the Integrated AUC of biomarkers selected by each metric in Table 1a. For comparison, the mean of the AUCs of individual genes selected by each metric was also calculated. However, without considering the diagnostic effects of genes together, the performance of the mean individual AUC is lower, as it cannot effectively capture the interactions between genes and is also more susceptible to outliers. In general, Integrated AUC has higher performance and is more robust, and will be used as the main performance measure in this study.

The top local and global metrics found are Clustering Coefficient (AUC=0.8770) and Bottleneck (AUC=0.8609) respectively (Table 2). Compared to the commonly used Degree, the AUC of Clustering Coefficient is 17% higher. Rather than simply using Degree to select genes with many connections, these results explore and confirm that other scoring methods better capture the functional features of protein networks. With Integrated AUC, we can validate the connection between the geometric metrics and their functional significance.

**Table 2.** Scoring metrics ranked by AUC

| Scoring metric | Integrated AUC | Mean Individual AUC |
| --- | --- | --- |
| **Clustering Coefficient** | 0.8770 | 0.8159 |
| **Bottleneck** | 0.8609 | 0.8012 |
| **Betweenness** | 0.8517 | 0.8091 |
| **Eccentricity** | 0.8409 | 0.8017 |
| **Stress** | 0.8179 | 0.7919 |
| **DMNC** | 0.8136 | 0.7860 |
| **MCC** | 0.8114 | 0.7874 |
| **EPC** | 0.8030 | 0.7959 |
| **Degree** | 0.7875 | 0.7877 |
| **MNC** | 0.7748 | 0.7890 |
| **Radiality** | 0.7904 | 0.8100 |
| **Closeness** | 0.6366 | 0.7986 |

#### Design of C-Index

In this second stage, we systematically composed metrics to find an ideal set of scoring methods that can adequately capture the interactions in biological networks without incurring high complexity. We evaluated composite performances using Integrated AUC of the top 10 and 20 genes of the superset of the metrics, and their Precision in correctly identifying the top 10 and 20 overall biomarkers found in Table 1b.

To measure both the node-level properties and the network-level properties, we chose to compose the top local and global metrics (which were also the top two metrics overall), Clustering Coefficient and Bottleneck (Figure 5). 7 out of the top 10 overall genes were correctly identified, resulting in a 70% precision, and the Integrated AUC of the top 10 nodes was 0.9027. Likewise, the top 20 nodes had 70% precision, and Integrated AUC was 0.8765. Already, the composition results in a higher AUC than any of the individual metrics.

**Figure 5.**
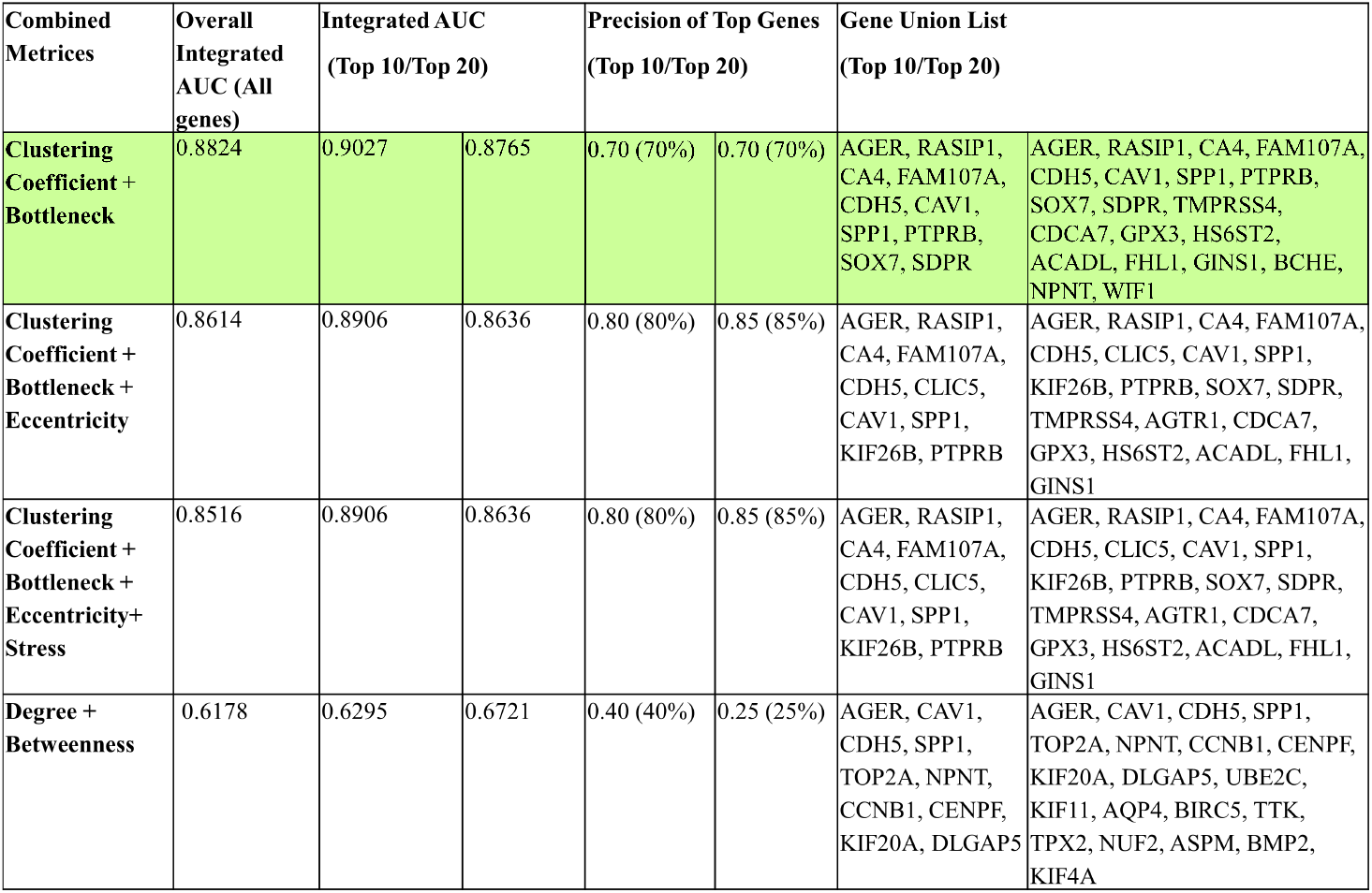
Performance of composite network scoring metrics

To determine whether the composition of Clustering Coefficient and Bottleneck is the most optimal, we also composed the three metrics Clustering Coefficient, Bottleneck, and Eccentricity. Although Betweenness outperformed Eccentricity, we chose to use Eccentricity because by definition Betweenness is essentially the same measure as Bottleneck and is thus not needed in the composite. With three, although the precision increased, the Integrated AUC reduced slightly. We also composed the top four metrics but obtained no improvement in AUC or precision. Considering the vastly increased complexity of adding more metrics, using two metrics together is ultimately more powerful than using three or more.

Our investigations have shown that Clustering Coefficient and Bottleneck can work together to adequately capture the significant biomarkers in the network with the most efficiency, and we propose to compose them as a new metric, **C-Index**. It outperforms all individual metrics and other compositions. It can achieve the comprehensiveness of using all metrics while vastly decreasing the complexity and inefficiency. Clustering Coefficient and Bottleneck work together as local and global metrics to better capture the nature of cancer and are capable of identifying essential genes that are prevalent throughout the entire network as well as locally.

Most literature studies use Degree and occasionally Betweenness to find their top biomarkers^23^. To investigate whether the combination of these two metrics results in effective biomarker selection, we calculated the performance of their composition. We obtained shockingly poor results: the top 10 genes had 40% precision, and 0.6295 Integrated AUC, while the top 20 genes had only 25% precision, and 0.6721 Integrated AUC. This poor performance is due to several factors that are often overlooked by other studies. Degree, a local metric, only measures how many connections a node has, which may signify that it regulates many other genes. However, these highly connected “hubs” have no great overall influence in cancer-causing pathways because their neighbors are not necessarily interconnected. In contrast, the Clustering Coefficient of a network, also a local metric, measures the tendency of nodes to form densely connected communities. In biological networks, these communities signify functional modules and gene complexes that work closely together and share similar functions. Clustering Coefficient identifies key biomarkers located in significant communities that work together to induce cancer and is a much more robust local metric than Degree in locating essential genes.

Although Clustering Coefficient alone is a powerful metric, it is local and unable to capture the overall topologies of the network. To gain a better view of the essential genes that play important roles in the network as a whole, Bottleneck is used complementary with Clustering Coefficient. Bottlenecks, genes with the highest betweenness centrality, control most of the information flow and interaction between proteins and are key connectors between regulatory pathways that are identified through Clustering Coefficient.

Together, Clustering Coefficient and Bottleneck are capable of considering multiple biological pathways, and improve on the performance of Degree from 78.75% to 88.24%. Not only does C-Index perform better than individual metrics, but it matches the performance of using all 12 metrics together with greatly reduced complexity and cost. The AUC of the overall top 10 biomarkers identified using all 12 metrics was 0.9064, as calculated in Stage I of this study. The AUC of the top 10 biomarkers identified using C-Index was 0.9027. It is as powerful as using all twelve, while much more efficient.

Our results indicate that the proposed C-Index is capable of capturing the most critical group of interactive genes that can early diagnose cancer. Our method of evaluating scoring methods is transformative and a breakthrough in applying topological analysis to biomarker identification.

#### Performance of C-Index in validation set

To validate the C-Index, we calculated the Integrated AUC of the metrics in the 20% validation set. As predicted, C-index outperformed Degree and each of its components (Supplementary Table 2. Compared to Degree, the performance of using C-Index to select biomarkers increased by 25%. Clustering Coefficient alone improved on Degree by 22.4%. Compared to using all 82 genes found by all 12 metrics, the performance of using only genes found by C-Index is 40% higher. This demonstrates that the metrics that compose C-Index vastly improve on conventional ones, and are capable of identifying a concise list of significant genes most successfully. Benefiting from both metrics with close interactions locally and genes that lie on the critical paths linked globally, C-index can revolutionarily select the most representative group of biomarkers at a low computational cost for accurate diagnosis.

#### Machine learning validation of biomarkers and metrics

We further validated the above results with a machine learning model we designed, the multi-stage Cascading Model (Methods). The model considers not only the genetic attributes, but also clinical attributes such as age, gender, and smoking status that most impact the development of NSCLC to improve the accuracy of diagnosis. Literature studies often decouple genomic and clinical attributes in cancer prediction. To incorporate both types of attributes and provide a validation method that does not involve Integrated AUC, we design this clinicogenomic model to explore their potential for disease diagnosis.

With the addition of the biomarkers, the purely clinical model improved by 51% (Fig.6a). Compared to using biomarkers alone, the addition of clinical variates also increased performance. The 4-gene model once again outperforms the top 10 model, validating the effectiveness of the signature. The 4-gene model was further extended to diagnose early vs. late disease stage, as well as NSCLC stages I-IV (Fig.6c).

**Figure 6.**
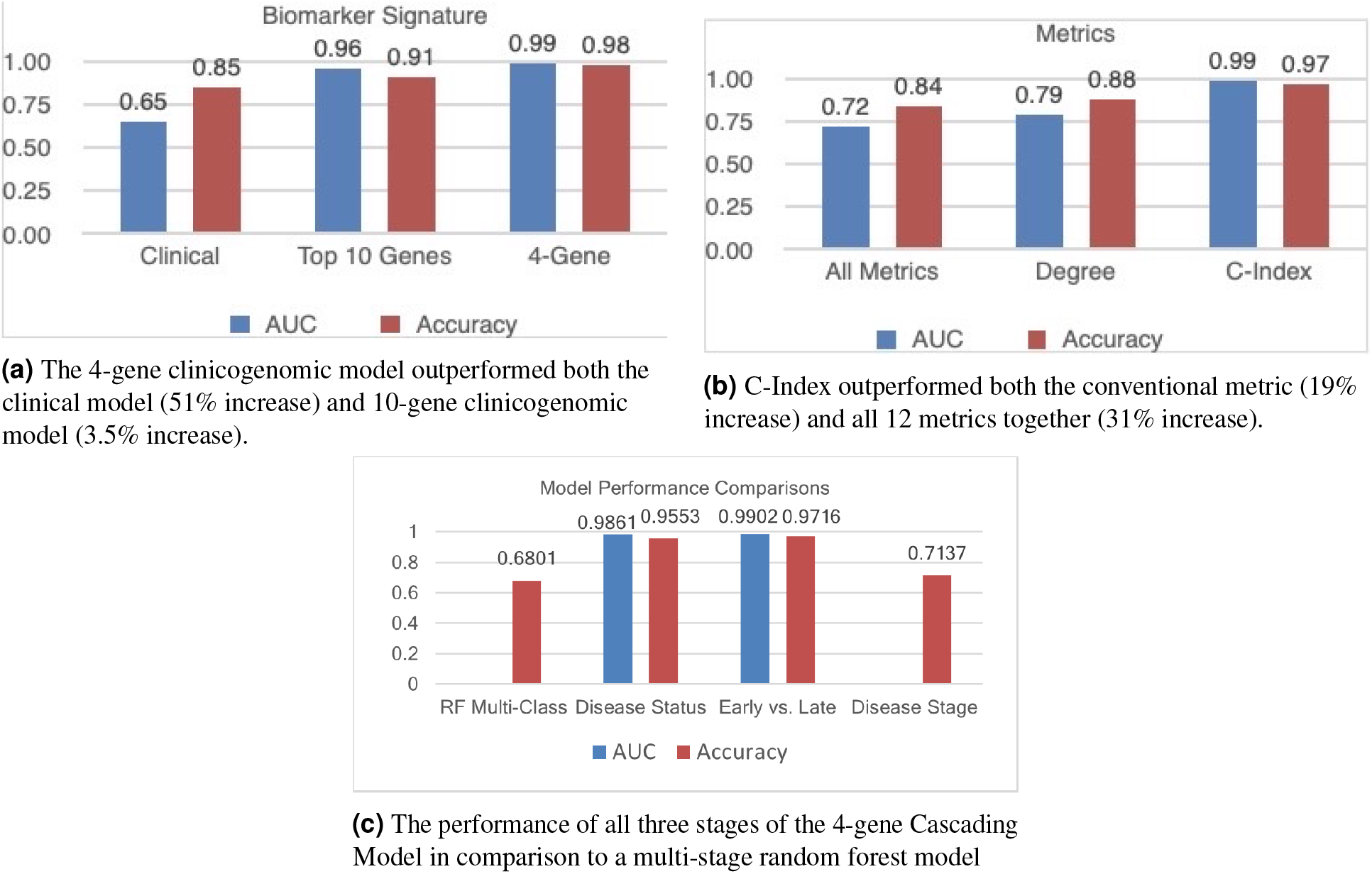
Machine learning validation of biomarkers and metrics

C-Index outperformed the conventional metric by 19% and all the metrics together by 31% (Fig. 6b). Utilizing all metrics suffered from the existence of outliers and inaccuracy. Degree, on the other side of the spectrum, was unable to identify a complete set of genes that could accurately capture the interactions of the network and thus performed poorly.

## Discussion

It is vitally important to identify critical biomarkers for exploring the pathogenesis of NSCLC, one of the deadliest cancers in the world. The key to improving prevention and early diagnosis is to find metrics and methods that can guide the effective yet cost efficient search of biomarkers.

There are several important findings from this study. First, through a series of functional, network, and statistical analyses, we identify a 4-gene biomarker signature consisting of *AGER, CA4, RASIP1*, and *CAV1* that can be further explored as possible therapeutic targets for drug treatment. Second, we prove that the most widely used topological scoring metric, Degree, is not the best suited for biological networks. We instead propose C-Index, a novel composite index that combines Clustering Coefficient and Bottleneck to best capture the interactions in gene networks for high efficiency and performance. Our results solidify the connection between geometric connectivity and functional connectivity. For validating the C-Index and 4-gene signature, we exploited the use of a machine learning model that considers both genomic and clinical factors concurrently.

This study largely focused on biological implications. Through functional enrichment analysis, we found that the four genes in the signature are enriched in GO terms of receptor activity, immune response, extracellular matrix, and signaling activity, which all play an important role in regulating proliferation, differentiation, and apoptosis. The significant downregulation of these four anti-tumor genes in cancerous patients signifies that a change in their expressions disrupts the tumor microenvironment, promoting tumorigenesis. These results match the significant GO and KEGG pathways identified in the first part of work. The significant biomarkers discovered may be further explored through knock-out trials and analysis of their impacts on cancer prognosis. Results from these trials can be applied to developing drug therapies that target these biomarkers.

The two metrics that compose C-Index, Clustering Coefficient and Bottleneck, effectively capture local and global gene interactions in the network. We hypothesize that Clustering Coefficient identifies significant communities that are most likely involved in pathways to promote tumorigenesis. Bottlenecks are critical points in the network that connect the biological pathways identified through Clustering. These two metrics work in tandem to effectively identify genes that work together in pathways to induce cancer.

Our research has shown that using multiple biomarkers and methods concurrently greatly improves performance. This is because genes work together in pathways that lead to tumorigenesis, and single genes cannot cause cancer without having numerous regulatory effects on other genes through signal transduction pathways. Cancer is a complex disease caused by the interaction of multiple environmental factors and genes. It is the combined effect of all these genes in the pathway together that leads to cancer onset. The 4-gene biomarker signature and biomarkers selected by C-index accurately capture the nature of cancer. With further validation and refinement in other cancer datasets, they are promising for the study of biomarkers and all cancers. The C-Index greatly increases the efficiency and accuracy of future biomarker searches, allowing for the low-cost identification of biomarkers with great diagnostic capability. The findings from this study provide an experimental foundation for further exploration of the usage of PPI networks to diagnose cancers. Most importantly, our results indicate that the conventional method of approaching TDA in oncology is greatly ineffective.

Topological analysis is a powerful method to analyze biomedical data. Our work lends itself to further exploration in settings other than biomarkers. Possible future directions include predicting treatment responses, cellular architecture determination, tumor segmentation, and other applications of cancer data. Our research can be expanded to other types of cancers and more datasets to validate our results and C-Index. The proposed methodology of finding top metrics can be extended to effectively and efficiently select biomarkers in various types of cancers, not just NSCLC, which helps to fundamentally advance the topological network research for the continuous pursuit of cancer prevention.

## Methods

### Dataset

In total, we analyzed 526 NSCLC samples consisting of 446 tumor and 80 control cases from the Gene Expression Omnibus (GEO) database^24^, a national repository of genetic information databases. In this study, we retrieved and combined three gene expression profiles (GSE31210, GSE33356, and GSE50081) to ensure greater accuracy and comprehensiveness. Finally, the merged dataset was divided into 80% for identification of top biomarkers and metrics, and 20% for validation.

### Data preprocessing and DEG identification

GEO2R analysis was performed to detect DEGs in NSCLC cancer tissues compared with normal samples. An initial pool of 267 statistically significant DEGs (p-value<0.01 and |logFC| > 1.2) were identified for further analysis and classified as up or down regulated. The raw gene expression values are normalized using z-score.

### GO and KEGG pathway enrichment analysis

Using the DAVID functional annotation tool, we performed enrichment analyses to identify significant GO terms with FDR < 0.05. The Database for Annotation, Visualization and Integrated Discovery (DAVID) gene functional annotation tool^25,26^ was used to identify significant GO (gene ontology) terms, which are biological annotations that signify functional characteristics. They are divided into three main categories: molecular function (MF), cell composition (CC), and biological processes (BP). The most significant GO terms were analyzed using DAVID to identify enriched terms with a threshold value of FDR (adjusted p-value) < 0.05. Similar in function to the adjusted-p-value, the lower the FDR, the more significant the enrichment. Statistically significant GO terms were also expressed as a z-score expression

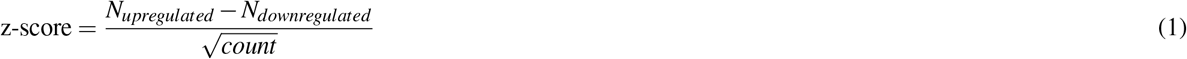

where *N*_*upregulated*_ and *N*_*downregulated*_ represent the number of upregulated and downregulated genes respectively. This expression value signifies whether the GO term is more likely to be downregulated (negative value) or upregulated (positive value). We visualized the top 10 GO terms and their z-score expressions using the GOplot package (version 1.0.2) in R^27^.

String-db^28^analysis was also utilized to identify significant KEGG pathways. The Kyoto Encyclopedia of Genes and Genomes (KEGG) database contains genomic information, functions, recognized pathways, and networks with higher-order functional information of various organisms^29^.

### PPI network construction

We constructed a protein-protein interaction (PPI) network based on the DEGs using the STRING Interactome database^28^, then inputted the network into *Cytoscape*^21^ for further analysis with the CytoHubba^22^ plugin. Using all 12 topological analysis methods, we identified our list of candidate biomarkers composed of the top genes selected by each method.

### AUC performance evaluation and Integrated AUC

We evaluated the diagnostic value and functional significance of the biomarkers using AUC, which measures the trade-off between sensitivity and specificity. AUC was calculated for each gene using the pROC package in R Studio^30^, which was also used to visualize the ROC curves. Similarly, to evaluate the performance of multiple biomarkers at once, Integrated AUC was calculated by first aggregating (in this study, we use mean) the expression values of the biomarkers, and then evaluating the AUC of that aggregated expression. To identify our 4-gene signature, we systematically aggregated the top biomarker expression values one by one and determined when the AUC reaches its peak.

### Survival analysis and validation of biomarkers

The Kaplan–Meier plotter database (http://kmplot.com) is an online tool used to analyze the associations between the identified hub genes and overall survival. The overall survival (OS) plots were based on 1925 lung cancer patients from GEO and TGCA (The Cancer Genome Atlas database) datasets. The hazard ratio (HR) with 95% confidence intervals and log rank P-value were calculated, and p-value < 0.05 was used to indicate a statistically significant difference.

The expression levels of the hub genes and their association with survival were additionally assessed using the web-based GEPIA database (http://gepia.cancer-pku.cn/)^31^ with the settings of p<0.05 and |Log2FC|>1.

### Evaluation of composite metrics

To evaluate composite metrics, we first fused, or supserset, the genes identified through each individual metric. Then, Integrated AUC of the top 10 and 20 genes of the superset was evaluated to ensure a fair assessment of each composition and to eliminate any bias surrounding using more or less number of biomarkers.

In addition to AUC, to quantify the ability of each metric to correctly identify the biomarkers that were included in the overall top 10 and top 20 ranked biomarkers found in Table 1, we also define a Precision metric as follows:

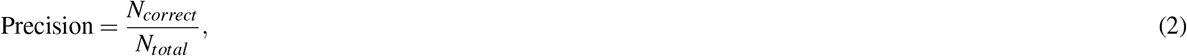

where *N*_*correct*_ and *N*_*total*_ represent the number of correctly identified genes and total number of top ranked genes respectively.

To find the number of correctly identified genes, we took the top 10 and 20 genes from the supersets of metric-selected genes and checked how many of them matched with the overall top 10 and 20 in the dataset.

### Cascading Model design

In addition to validating our results in the 20% validation set, we designed a Cascading machine learning model that evaluates the ability of biomarkers and metrics to diagnose disease. The studies above focused on the performance of biomarkers and how to improve the search for them. However, compared to only using genetic information, the usage of a variety of other factors will help improve the accuracy of diagnosis. Some clinical attributes that impact the development of NSCLC are age, gender, and smoking status. In order to incorporate both clinical and genomic attributes, we explored the use of machine learning techniques to adequately consider multiple factors at once in disease prediction. Our aim is to create an accurate clinicogenomic diagnosis model that can exploit top biomarkers selected to accurately predict disease status, differentiate between early vs. late stage of cancer, and identify lung cancer stages I-IV.

As an initial study, we tested a simple multi-class Random Forest (RF) algorithm as it is considered to have an advantage over other classification algorithms in terms of robustness to overfitting, ability to handle non-linear data, and stability in the presence of outliers. However, this model resulting in low accuracy for classification of disease stages.

To more accurately identify the state of cancer, we propose a multi-stage Cascading Model with 3 stages, depicted in Supplementary Figure 1. The model was used to validate the metrics and biomarkers in this study, and their capability of diagnosis. The model first classifies the data into disease or no disease, then further classifies those with disease into early vs. late stages of cancer, and finally classifies early and late one step further into cancer stages I-IV.

In the 4-gene model, the first classification disease status had an average accuracy of 0.9553 and AUC of 0.98605. The second classification, early vs. late stages, had an average accuracy of 0.9716 and AUC of 0.9902. Disease stages I-IV classification had an average accuracy of 0.7137, which may be due to the minor difference between the four disease stages of patients. The difference of clinical attributes between cancer and no cancer and early vs. late is a lot greater than the difference between the four stages. The model may have difficulty distinguishing the two sides of the boundary.

As exemplified in the performance study, the model greatly improves the ability to accurately differentiate between multiple stages of disease, and is one of the first capable of accurately predicting early vs. late cancer stages.

## Data Availability

The datasets GSE31210, GSE33356, and GSE50081 are available online from the GEO database.

## Code Availability

https://github.com/iwu24/NSCLCcascadingmodel.git

## Acknowledgements

We would like to thank Siya Goel from Stanford University for many helpful discussions throughout this613 wor

## Author contributions statement

I.W. conceived and conducted the experiment and analyzed the data. I.W. and X.W. wrote the paper. All authors reviewed the manuscript.

